# Endophytic colonization pathways of *Pseudomonas chlororaphis* M71 and *Trichoderma atroviride* SC1 in grapevine following stem injection

**DOI:** 10.64898/2026.07.10.737717

**Authors:** Greta Brussi, Alessio Martini, Claudio Ratti, Gerardo Puopolo, Laura Mugnai, Ilaria Pertot

## Abstract

Endophytic biocontrol agents may contribute to grapevine health, but their ability to establish, persist, and move within woody tissues remains poorly understood. In this study, a stem injection method was developed to introduce *Pseudomonas chlororaphis* M71 and *Trichoderma atroviride* SC1 into rooted and grafted grapevine plants, and their spatial and temporal colonization patterns were compared with the movement of a dye tracer. The dye tracer moved rapidly through xylem tissues, whereas both microorganisms showed more restricted early distribution. Over time, M71 and SC1 displayed distinct colonization patterns. M71 persisted after injection, but remained localized near the inoculation site, with limited movement toward roots or distal aerial tissues. In grafted plants, M71 recovery depended on the injection site and declined more markedly after rootstock injection than after scion injection. In contrast, SC1 showed broader and more persistent colonization. In rooted cuttings, SC1 was recovered from stem and root tissues up to 56 days post-injection, and in grafted plants it was recovered across the graft union, particularly after scion injection. Microscopy supported internal localization of both microorganisms. GFP-labelled M71 and SC1 hyphae were observed mainly within xylem vessels, and viable microorganisms were recovered from corresponding wood tissues. No contamination was observed in control plants. These results show that beneficial microorganisms can be introduced into grapevine tissues by stem injection and that bacterial and fungal biocontrol agents differ markedly in their internal movement and persistence.

**IMPORTANCE:** Introducing beneficial microorganisms directly into plant tissues could help in establishing protective endophytic populations, but little is known about how such microorganisms move and persist inside grapevine. This study shows that stem injection can deliver *Pseudomonas chlororaphis* M71 and *Trichoderma atroviride* SC1 into grapevine tissues without visible phytotoxicity. The two microorganisms followed different colonization patterns. M71 remained mostly localized near the injection site, whereas SC1 spread more broadly and persisted in both rooted and grafted plants. These findings provide a basis for developing targeted endophytic biocontrol strategies in grapevine propagation and early plant establishment. This approach may be particularly relevant for grapevine trunk diseases and other vascular disorders, in which pathogens colonize internal woody tissues.

## INTRODUCTION

In natural environments, plants function as complex ecosystems in which interactions with rhizospheric, epiphytic and endophytic microbial communities form dynamic networks that influence plant development, defense, and responses to environmental stimuli (1). Among plant-associated microbes, endophytes, defined as microorganisms that spend at least parts of their life cycle inside plants (2), have received increasing attention due to their ecological functions and their potential application in agriculture (3; 4).

Endophytes can colonize internal plant tissues through multiple entry routes and establish in specific microbial niches, which are influenced by the specific organ involved (5). Due to the dense microbe-microbe and microbe-host interactions, the rhizosphere is the primary reservoir and recruitment zone for endophytes, from which microorganisms access internal plant tissues via root-associated entry sites. Consequently, this region is also a major source for the discovery of biocontrol agents (6, 7). The complexity of biological activity and chemical signaling in the rhizosphere poses significant constraints that endophytes must overcome to achieve a successful plant tissue colonization and a stable host association (8, 9). Aerial organs, such as leaves, flowers and stems, also act as sources of endophytes (10), due to the constant exposure to the microorganisms from the environment. Seeds and cuttings can contribute to the vertical transmission of endophytes (11), while insect feeding activity may further facilitate their dispersal within and among plants (12). Unlike phytopathogens, endophytes typically colonize and move within plant tissues without causing detectable damage (13). They normally exploit natural entry points, such as stomata (14), hydathodes, and wounds (2), although some endophytes are capable of secreting enzymes that promote localized cell lysis and facilitate tissue penetration (15).

Endophytes are increasingly recognized as key contributors to health, nutrition, and overall performance of plants, including *Vitis vinifera* (16). For example, they can enhance nutrient acquisition and promote root and shoot development (17). Grapevine endophytes also contribute to host tolerance to abiotic and biotic stresses by modulating plant defense responses and secondary metabolism (18).

Microbial biocontrol agents are increasingly used to promote plant health and suppress pathogens, through mechanisms that include antibiosis, competition for nutrients/space, production of lytic enzymes, and induction of plant defense responses (19). However, most studies focus on rhizosphere or foliar applications, and achieving long-term colonization of grapevine tissues by beneficial endophytes remains challenging, because populations often decline or disappear over time, especially after transferring inoculated plants to non-sterile environment or field conditions (20, 21).

Inoculating biocontrol agents directly into plant tissue could, in principle, enable long-term and stable interactions and possibly prevent several diseases on perennials as grapevine (4, 22, 23). If established as endophytes, these microorganisms could help restore microbial balance, enhance systemic resistance, and promote overall plant growth (24, 2, 25). However, a crucial first step is to determine whether the introduced microorganism can successfully establish and spread within plant tissues. Standard inoculation methods (e.g. root dipping, foliar sprays, soil drenching) often fail to ensure consistent internal colonization by biocontrol agents (26, 27, 28, 20). Successful colonization depends on multiple factors, such as host species, inoculation technique, microbial density, and competion with resident microbiota, therefore achieving consistent long-term colonization remains a major challenge for practical biocontrol applications (20, 28). Some attempts to manipulate endophytic microbiome have already been undertaken with various microbial inoculation strategies. For instance, injecting a consortium of selected endophytes altered the xylem microbiome in olive trees (29) and co-inoculating *Paraburkholderia phytofirmans* into grapevines with the pathogen via stem puncture reduced *Xylella fastidiosa* population size and related Pierce’s disease symptom severity (30).

*Pseudomonas* and *Trichoderma* species are among the most extensively studied bacterial and fungal biocontrol agents, respectively (31, 32, 33), although quantitative data on their movement, distribution, and persistence inside woody tissues are limited (34, 35). Various *Pseudomonas* species and strains have been identified as internal colonizers of grapevine (36), and several endophytic strains are being investigated for their potential to control grapevine diseases (37, 38). *Trichoderma* species effectively colonize inoculation sites, such as pruning wounds in grapevines, and can persist on the plant for several months after application (39, 40). Their presence as natural grapevine endophytes have also been documented (41, 34), and various strains currently used or tested as biocontrol agents were originally isolated from grapevine tissues (42, 43).

The aim of this study was to investigate the spatial and temporal colonization of two beneficial microorganisms after stem injection in grapevine tissues. Specifically, we developed a stem injection method and evaluated persistence and internal movement of a bacterial strain (*P. chlororaphis* M71) and of a fungal strain (*T. atroviride* SC1) in rooted and grafted grapevine plants.

## Materials and methods

### Microorganisms

*Pseudomonas chlororaphis* M71 (hereafter M71) was isolated from the rhizosphere of a tomato plant grown in a farm located in the municipality of Salerno (Italy) and carries a rifampicin resistance marker (100 µg mL^-1^) that enables its selective recovery from plant tissues (44). *Trichoderma atroviride* SC1 (hereafter SC1) was obtained from the culture collection of Fondazione Edmund Mach, San Michele all’Adige (Italy) and can be unambiguously identified according to the method of Savazzini et al. (2008) (45).

For bacterial inoculum preparation, M71 was cultured on Nutrient Agar (NA; Thermo Fisher, USA) in 90-mm Petri dishes at 27 °C for 48 h. Three colonies were harvested using a sterile inoculating loop and transferred into 100 mL of Luria-Bertani broth (LB; Sigma-Aldrich, USA) in 250 mL Erlenmeyer flasks and incubated at 27 °C for 24 h in an orbital shaker (Lab Companion Si-600R, Jeio Tech, Republic of Korea) at 180 rpm. Cultures (50 mL) were centrifuged at 5000 rpm for 10 min (SL 16R Centrifuge, Thermo Fisher) in sterile Falcon tubes and the resulting pellets were suspended in sterile physiological saline solution (SPSS; 0.85% NaCl). Cell density was adjusted to an optical density at 600 nm (OD₆₀₀) of 0.3, corresponding to approximately 1 × 10⁸ colony forming units (CFU) mL^-1^, as determined spectrophotometrically (Ultrospec 3100 pro, Amersham Biosciences, UK). Bacterial concentration and viability were confirmed by serial dilution and plate counting on NA supplemented with rifampicin.

SC1 was maintained on Potato Dextrose Agar (PDA; Thermo Fisher) in slants at 4°C until use. Inoculum was prepared by culturing SC1 on the same medium in 90-mm Petri dishes at 25°C until conidiation (two weeks). Conidia were harvested by flooding the colony surface with 2 mL of SPSS and gently scraping with a sterile loop. The final conidial concentration was determined using a Thoma counting chamber under a light microscope. The final concentration was adjusted to 1 × 10⁶ conidia mL⁻¹. Conidia viability was assessed by germination tests on PDA and only suspensions with germination rates ≥ 95% were used. Both bacterial and fungal inocula were freshly prepared prior to each experiment.

The presence of viable M71 and SC1 in the wood, leaf, and root samples of all treatments was assessed by colony growth on the respective selective (M71) or semi-selective (SC1) media. Namely, M71 was isolated on NA supplemented with 0.1 g L^-1^ rifampicin (Sigma-Aldrich) and 0.1 g L^-1^ cycloheximide (Sigma-Aldrich), while SC1 was recovered on PDA supplemented with 0.5 mg/L rose bengal (Thermo Fisher), 0.1 g/L chloramphenicol (Sigma-Aldrich) and 0.1 g L^-1^ streptomycin (Sigma-Aldrich). Petri dishes (90 mm diameter) were incubated at 27 °C for M71 and at 25 °C for SC1, until colonies were visible. Positive controls (suspensions of M71 and SC1) were included for both media. Because the semi-selective medium may also allow the growth of other *Trichoderma* species, the identity of SC1 was confirmed on a subsample of colonies (45).

### Plants and growth conditions

Dormant canes of *Vitis vinifera* L. cv. Pinot Noir (clone ENTAV 667) were collected from an experimental vineyard planted in 2018 at Fondazione Edmund Mach (46.1892°N, 11.1379°E). Canes were pruned on 15 December 2023 and stored at 7°C in a sealed black plastic bag until use. No chemical treatments were applied to the vineyard during 2023 to avoid any interference with original endogenous microbial communities and inoculated strains during the trials. On 15 January 2024, two-node cuttings (10-15 cm internode length) were prepared from stored canes and hydrated overnight in tap water at 22 ± 1°C. Cuttings were then planted in water-saturated sphagnum peat pellets (Jiffy-7 The original, Jiffy group, USA) and maintained in a greenhouse under controlled conditions (25 ± 1°C, 16 h light/8 h dark photoperiod, 70 ± 10% relative humidity) for the entire duration of the experiments. After 30 days, cuttings exhibiting uniform shoot growth, well-developed root system and intact stems were selected and transplanted into 1-L pots containing a commercial substrate (Vegetal Radic, TerComposti, Italy).

Grafted grapevine plants (cv. Pinot Noir, clone ENTAV 667, grafted onto SO4 rootstock; Vivai Cooperativi Rauscedo, Italy) were transplanted into the same type 1-L pots and maintained under the greenhouse conditions described above. To standardize vegetative growth, a single bud was retained per cutting/plant by removing all additional ones. All plants were irrigated as needed with tap water and were visually inspected to confirm the absence of symptoms associated with known grapevine diseases prior to inoculation.

### Stem injection

Preliminary trials were carried out to optimize injection depth and assess the solution movement within grapevine tissues using a dye tracer (safranin 0.5% w/v in distilled water, Sigma-Aldrich). Possible safranin phytotoxicity was excluded by collecting thin wood sections from dye-stained areas and examining xylem vessels under the light microscope for necrosis, cell wall degradation, and gum exudation. Based on the preliminary measurements (data not shown), a penetration depth of approximately 1 mm was set in order to reach the xylem vessels, whereas 2 mm were set to access the pith.

Stem injections on cuttings were carried out on the stem surface on the actively growing shoot’s side, a position selected because it favors upward translocation through the xylem and simulates natural sap flow. In rooted cuttings, the injection was performed on the stem of the cutting, approximately 5 cm below the base of the growing shoot. In grafted plants, the injection was performed either in the middle portion of the rootstock, below the graft union, or in the middle portion of the scion, above the graft union (Figure 1). Rooted cuttings and grafted plants were injected with the dye tracer, M71 and SC1 suspensions and the SPSS (negative control). Three plant (replicates) per treatment were used, with a total of 36 plants per trial. The trial was carried out twice under the same conditions.

**Figure 1.**
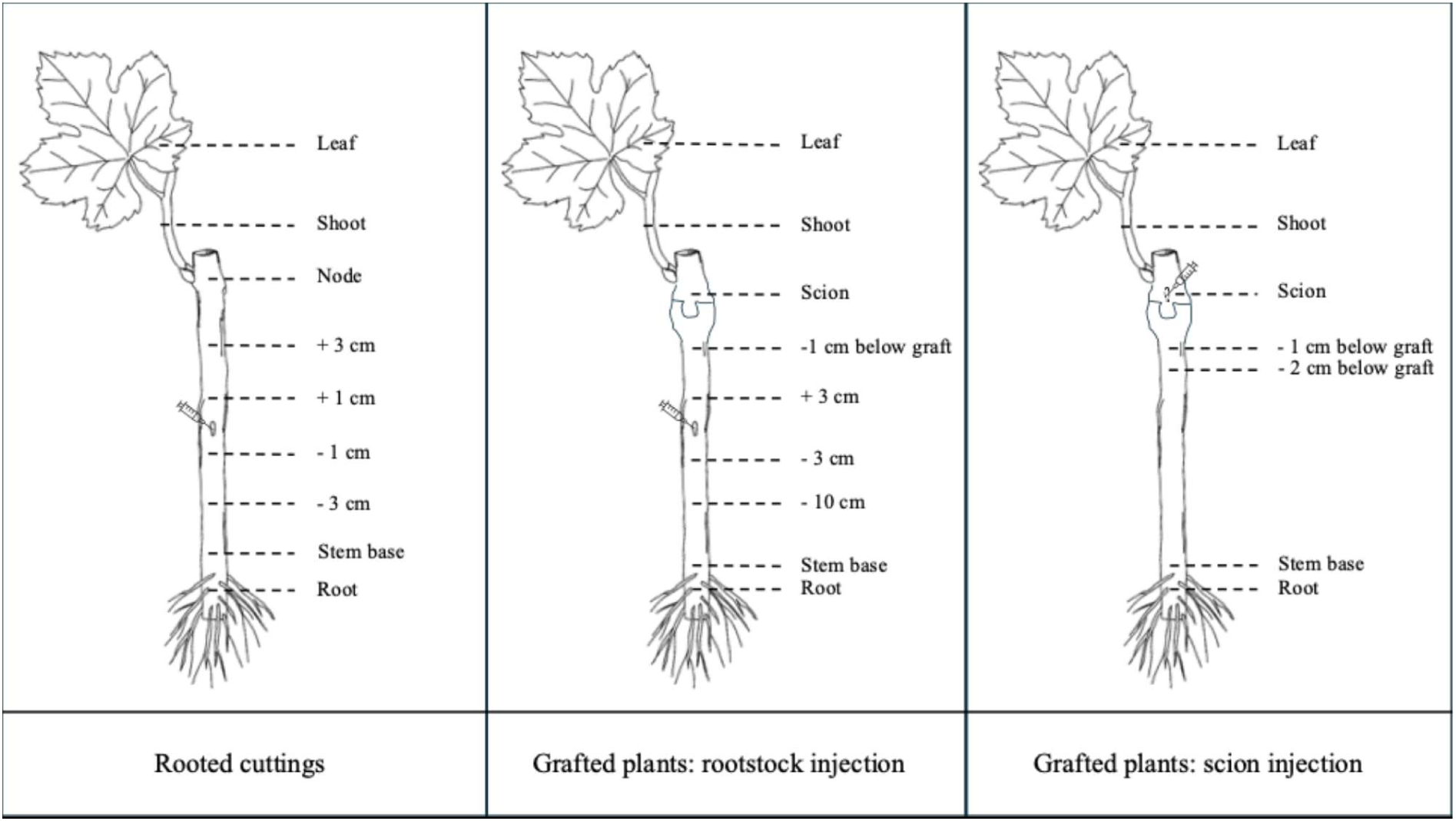
Schematic representation of the injection point and the sampling positions used to assess dye distribution and microorganism colonization 48 h after stem injection in grapevine plants. Distances are measured from the injection point.

The injection was carried out in all plants as follows. The stem surface was disinfected by spraying 70% ethanol. A sterile drill bit (Dremel 3000, Dremel, USA) was used to create a 1-mm diameter hole at a 45° upward angle. A sterile 1-mL pipette tip (Sarstedt, Germany), shortened by 2 mm at its narrow end, was inserted into the hole through a pre-perforated piece of parafilm. The parafilm was wrapped tightly around the stem to ensure sealing and to promote passive solution uptake. Using a sterile syringe, 1 mL of the dye solution was loaded into the pipette tip and allowed to enter the stem passively through capillary action. The solution level was marked on the pipette tip to monitor uptake over time and the volume of solution absorbed was approximately 0.3 mL. After 24 h, the pipette tip was removed and the injection site was sealed with parafilm.

To visualize and measure the distribution of the injected dye solution in the stem and root tissues, the bark was removed 48 h post injection. Dissection tools were flame-sterilized between sampling positions to avoid cross-contamination among tissues. In rooted cuttings, dye or microorganisms’ distribution was assessed at +1 and +3 cm above the injection site, and at -1 and -3 cm below the injection site, as well as at the stem base (2 cm above the root-stem junction), roots (at 2 cm below the root-stem junction), node (middle), shoot (1 cm from the base of the shoot), and leaf. In grafted plants that had been injected in the rootstock, dye or microorganisms’ distribution was assessed +3 cm above the injection site, -3 and -10 cm below the injection site, at the graft union region (below graft and scion), and in the stem base, roots, middle of shoot, and leaf. In grafted plants injected in the scion, dye or microorganisms’ distribution was assessed at the scion (middle), 1 and 2 cm below the graft union, and in the stem base, root, shoot, and leaf (Figure 1).

Distribution or colonization was assessed in terms of incidence and frequency. To standardize the assessment, each stem section was divided into four quadrants and dye distribution was recorded as the number of quadrants exhibiting visible red staining (range: 0-4). In the microorganism inoculated plants, the four quadrants (corresponding to approximately 0.5 g each) were aseptically excised and placed on the respective selective/semi-selective medium until exhibiting microbial growth. Color or colonization incidence was defined as the presence of red staining or microbial growth in at least one fragment per sampling position and was expressed as the percentage of each plant sampling position exhibiting colonization. Incidence indicates whether the dye or microorganism reached a given sampling position from the injection site and was used as an indirect measure of longitudinal distribution from the injection site. Color or colonization frequency was expressed for each sampling position as the percentage of tissue fragments showing red staining or yielding microbial growth. Frequency represents the proportion of tissue fragments or quadrants within each sampling position showing dye staining or microbial growth and therefore provides information on the extent of local or transverse spread/colonization within that section.

For root sampling the central cylinder of four randomly selected roots was collected. Leaf sampling was performed at defined positions, namely the petiole insertion site, the leaf tip, and the left and right medial lamina. Red staining was visually assessed on intact leaf tissue, while for microbial growth, leaf samples consisted of four 1-cm-diameter leaf discs collected with a sterile cork borer from the first fully expanded leaf at the node. Before taking the sampled tissues, all wood, root, and leaf samples were surface sterilized by 1% sodium hypochlorite, rinsed three times in sterile distilled water, and dried on sterile filter paper. Aliquots of the final rinse water were plated on the corresponding selective or semi-selective medium to verify the effectiveness of surface disinfection.

Quantification of the microorganisms in the tissues was carried out on additional rooted cuttings that were injected with M71 or SC1 (five cuttings for each treatment) and sampled at 28 dpi, 1 cm above and 1 cm below the injection site. After surface disinfection as described above, tissue samples (1 g) were ground in 10 mL SPSS in 50 mL Falcon tubes and agitated on the shaker at room temperature (25 °C) for 4 hours. Serial dilutions (from 10⁻¹ to 10⁻⁷) were plated. For M71, three 10 µL drops of each dilution (technical replicates) were plated on the selective medium in Petri dishes and incubated at 27 °C until colonies became visible. For SC1, three 100 µL aliquots were plated on the semi-selective medium in single Petri dishes (technical replicates), spread with an L-shaped scraper and incubated at 25°C. Colonies of M71 and SC1 were counted, and the mean of three technical replicates was used to calculate colony forming units per gram of fresh tissue (CFU g⁻¹) after adjusting for dilution factor and extraction volume (10 mL per 1 g tissue).

### Spatial-temporal colonization of *Pseudomonas chlororaphis* M71 and *Trichoderma atroviride* SC1 in grapevine rooted cuttings

To assess the colonization of the microorganisms over time within the grapevine tissues, rooted cuttings were injected 5 cm below the base of the growing shoot with M71, SC1 and SPSS, as described above. The presence of the inoculated microorganisms was assessed at 14, 28, and 56 days post-injection (dpi) at the sampling positions described above. Each treatment and time point consisted of seven replicates, with a total of 63 plants. The trial was carried out twice under the same conditions.

### Spatial-temporal colonization of *Pseudomonas chlororaphis* M71 and *Trichoderma atroviride* SC1 in grapevine grafted plants

To understand whether grafting position may influence the colonization of the microorganisms over time within grafted plants M71, SC1 and SPSS were injected in the middle of the rootstock or in the middle of the scion and sampled at 28 and 56 dpi with the same methodology described above. Each treatment and time point consisted of seven replicates, with a total of 84 plants. The trial was carried out twice under the same conditions.

### In planta visualization of Pseudomonas chlororaphis M71

To visualize the presence of M71 in grapevine tissues following injection, the strain was fluorescently labelled by chromosomal insertion of a green fluorescent protein (GFP) reporter using a triparental mating approach. Plasmid transfer was obtained by the helper strain *Escherichia coli* HB101 carrying plasmid pRK2013, which provides the mobilization functions (*mob* gene) required for the transfer. The donor strain was *E. coli* DH5α carrying pUT-miniTn5-Km and the recipient strain was M71. Donor and helper strains were grown overnight in LB broth at 37 °C with shaking (180 rpm), supplemented with kanamycin (Sigma-Aldrich) (50 µg mL⁻¹) and rifampicin (100 µg mL⁻¹), respectively. The recipient strain was cultured overnight at 27 °C with shaking (180 rpm) in LB broth containing rifampicin (100 µg mL⁻¹). The cultures (5 mL) of each strain were harvested by centrifugation (5,000 rpm, 5 min), washed three times with SPSS and resuspended in the same solution (5 mL). Donor, helper, and recipient strains were mixed in a volumetric ratio of 20:20:1 (final volume of 1 mL). Typically, 488 µL of donor, 488 µL of helper, and 24.4 µL of recipient were combined. Aliquots (100 µL) of the mixture were spotted onto LB agar in Petri dishes without antibiotics and incubated at 29±1 °C for 24–48 h until visible colony growth. Donor-only and recipient-only controls were included. After incubation, bacterial spots were resuspended in 1 mL SPSS. Aliquots of 100 µL were plated onto selective LB agar (added with kanamycin 50 µg mL⁻¹ and rifampicin 100 µg mL⁻¹) to recover transconjugants. Petri dishes were incubated at 27°C and colonies were checked daily between 24-72 h. Successful plasmid mobilization was confirmed by the absence of colonies of the donor and recipient strains on selective medium supplemented with both the kanamycin and rifampicin.

To verify that the fluorescent labelling did not alter the biological characteristics of M71, the transformed strain was compared with the wild-type in terms of growth kinetics, colony morphology and production of phenazines, evidenced by the presence of the characteristic orange halo on agar medium. Biocontrol activity was evaluated using dual culture assays against *Phaeomoniella chlamydospora* and *Neofusicoccum parvum* that served as reference organisms.

The GFP-labelled M71 was injected in six rooted cuttings, following the same methodology described above. Six additional cuttings were injected with SPSS (negative control). At 28 dpi, six transverse wood slices (1.5 cm) were obtained from three cuttings at 2 cm above and 2 cm below the injection site using a cryostat and observed under a fluorescence microscope (Eclipse 80i, Nikon, Japan). GFP fluorescence was detected using excitation/emission filter sets 420–490 nm (B-3A) and 450–490 nm (B-2A), while plant tissue autofluorescence was assessed using a DAPI filter (340–380 nm). Observations were done in xylem and phloem vessels, adjacent parenchyma tissues and pith. Corresponding wood sections of M71 injected rooted cuttings were collected and incubated in Petri dishes containing NA supplemented with rifampicin at 27 °C for five days to assess the viability of the transformed M71 strain and verify the presence of fluorescence under the same microscope setting in the resulting colonies. Corresponding wood sections of SPSS injected rooted cuttings were incubated on the same medium to verify the absence of M71.

### In planta *visualization of* Trichoderma atroviride *SC1*

To visualize the presence of SC1 in the grapevine tissues following injection, six rooted cuttings were injected as described above and processed 28 dpi at the same positions used for microbial isolation in the rooted cuttings experiment. Five cuttings injected with SPSS were added as negative controls. The stem of the rooted cutting was cut into approximately 1.5 cm-long segments, each segment was split lengthwise into four sections, and the bark was removed. Then, sample preparation followed the methodology of Proto et al. (2022) (46). Briefly, samples were fixed in 5% (v/v) glutaraldehyde (Merck KGaA, Germany) in 0.1 M phosphate buffer solution (pH 7.4) at 4 °C for 48 h to fix tissues, rinsed in the same buffer for 5 minutes and dehydrated in a graded ethanol series (20, 30, 50, 75, 95 and 100%). Dehydrated samples were then critical-point dried with liquid CO_2_ (CPD K850, Electron Microscopy Sciences, USA) for 2 h, sputtered-coated with silver (K500 sputter, Emitech, USA) and observed under SEM (SEM 515, Philips, Netherlands) operating at 30 kV. All tissue types were observed, with particular attention to the vascular tissue and the pith. Corresponding wood sections were collected and incubated in Petri dishes containing the semi-selective medium to verify the presence of SC1. Corresponding wood sections of SPSS injected rooted cutting were incubated on the same medium to verify the absence of SC1.

### Data collection and statistical analysis

To evaluate the relationship between dye distribution and isolation of the two microorganisms at 48 h post injection, both color or colonization incidence and color or colonization frequency were compared. Color or colonization incidence was compared between dye and microorganisms using counts of presence/absence per position using Fisher’s exact test with Monte Carlo for each experimental setup (rooted cuttings, grafted plants injected in the rootstock and grafted plants injected in the scion). Color or colonization frequency (percentage of colored or colonized tissue fragments per position) was assessed using Spearman’s rank correlation coefficient (ρ) between dye and microorganism patterns.

Regarding microorganism isolation in rooted cuttings and grafted plants at the various time points post injection, colonization incidence was analyzed using generalized linear mixed model (GLMM) with binomial distribution and logit link function, including position as fixed effect and plant as random effect. Colonization frequency was analyzed as counts per position using GLMMs with Poisson distribution, with the same setting. Post-hoc pairwise comparisons between positions were performed using Tukey’s Honest Significant Difference (HSD) test (α = 0.05).

For each experiment, data from the two repeated trials were pooled for analysis after confirming the absence of significant differences between trials using Fisher’s exact test for colonization incidence data and Mann-Whitney U test for colonization frequency data. All the analyses were conducted in R (version 4.4.0).

## Results

### Stem injection

To evaluate the movement of the injected suspensions within grapevine tissues, the distribution of the dye tracer was compared with the colonization patterns of M71 and SC1 48 h after stem injection. No M71 or SC1 colonies were recovered from plants injected with the dye tracer or with SPSS, confirming the absence of cross-contamination and the specificity of the microbial detection protocol. No phytotoxicity symptoms were observed in either dye tracer-injected or microorganism-injected plants.

Overall, after 48 h, the dye tracer was detected over a broader longitudinal range than either microorganism (Figure 2). By contrast, M71 and SC1 colonization was mainly restricted to tissues close to the injection site.

**Figure 2.**
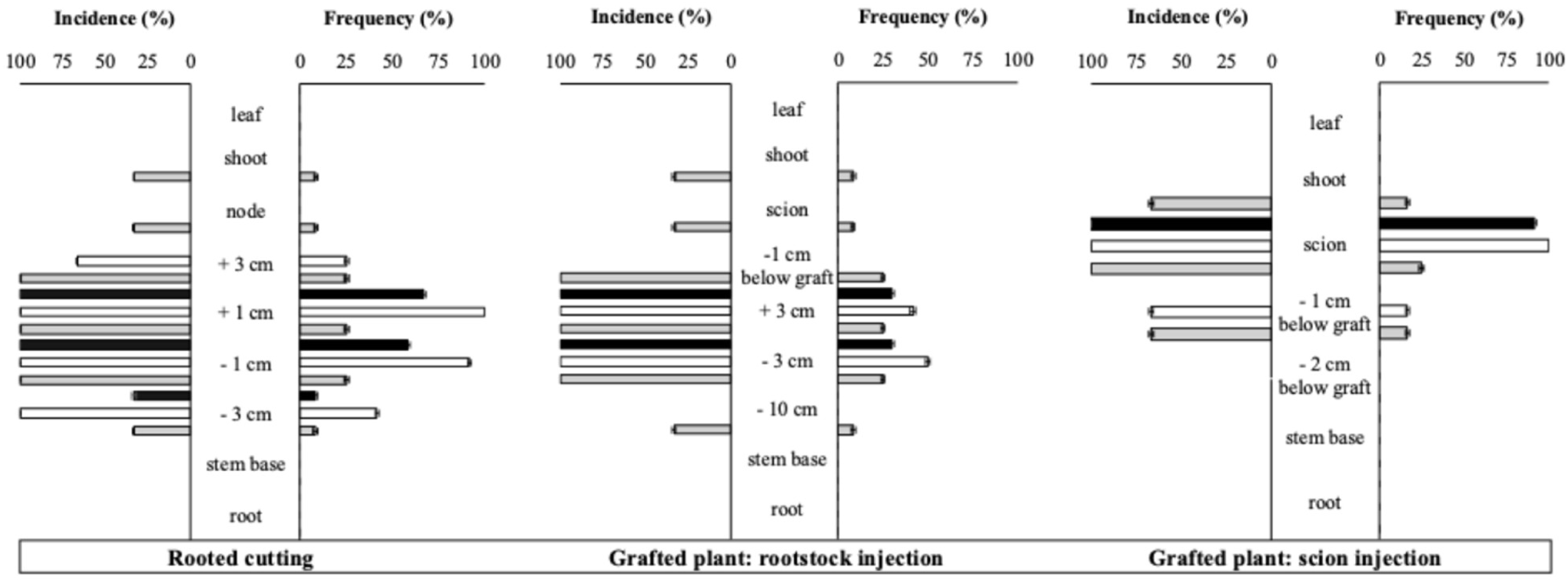
Incidence and frequency of dye distribution (grey histogram), Pseudomonas chlororaphis M71 (black histogram), and Trichoderma atroviride SC1 presence (white histogram), 48h after injection. Assessments were carried out in rooted cutting injected in the middle of the stem and grafted plant injected in the middle of the rootstock or in the middle of the scion. Error bar indicate standard error.

In rooted cuttings, the dye was detected primarily above and below the injection point, particularly at +3, +1, and -1 cm, whereas little or no staining was observed in distal tissues such as the stem base, roots, or leaves. M71 and SC1 showed a broadly similar but more localized distribution. M71 was mainly recovered near the injection point, particularly at +1 and -1 cm, while SC1 was also detected at relatively high frequency at +3 cm.

A similar pattern was observed in grafted plants injected in the rootstock. Dye staining was concentrated around the injection region, especially at +3 and -3 cm and near the graft union, with lower incidence in the node and shoot. Microbial recovery was more restricted, with both M71 and SC1 detected only at positions close to the injection point, mainly +3 and -3 cm.

In grafted plants injected in the scion, dye staining was observed in the scion, in the shoot, and at -1 cm below the graft region. M71 and SC1 were recovered mainly from the scion close to the injection site; SC1 was also detected at -1 cm below the graft region. Neither microorganism was detected from -2 cm below the graft union nor in the shoot or leaf. Dye incidence was high across longitudinal sampling positions, whereas dye frequency within individual stem sections was lower. Staining was visually confined mainly to xylem vessels adjacent to the injection pathway, whereas microbial recovery, when present, was detected across a wider proportion of the sampled fragments.

The distribution patterns of the dye tracer, M71, and SC1 were compared using binomial generalized linear mixed models, with treatment and sampling position included as fixed effects, and plant identity included as a random effect. In rooted cuttings, incidence did not differ significantly among treatments, although a marginal treatment effect was observed (Wald χ² = 4.81, df = 2, *P* = 0.090). Pairwise comparisons showed no difference between dye and SC1, while dye and M71 tended to differ, but not significantly after FDR correction (dye vs M71, P = 0.069; dye vs SC1, P = 1.000; M71 vs SC1, P = 0.069). Sampling position also had only a marginal effect on incidence (Wald χ² = 4.81, df = 2, P = 0.090). In contrast, frequency differed significantly among treatments (Wald χ² = 19.20, df = 2, P < 0.001) and sampling positions (Wald χ² = 35.55, df = 4, P < 0.001). SC1 showed significantly higher frequency than both dye and M71 (dye vs SC1, P < 0.0001; M71 vs SC1, P = 0.0016), whereas dye and M71 did not differ significantly (P = 0.208). Thus, in rooted cuttings, SC1 showed greater local spread within sampled stem sections, while incidence patterns were broadly comparable among treatments. In grafted plants injected in the rootstock, incidence differed significantly among treatments (Wald χ² = 6.31, df = 2, P = 0.043), but not among sampling positions (Wald χ² = 6.46, df = 5, P = 0.264). M71 showed significantly lower incidence than both the dye tracer and SC1 (dye vs M71, P = 0.031; M71 vs SC1, P = 0.031), whereas dye and SC1 did not differ significantly (P = 0.071). With regard to frequency, no significant treatment effect was detected (Wald χ² = 4.03, df = 2, P = 0.133), and all pairwise comparisons among treatments were non-significant. However, frequency differed significantly among sampling positions (Wald χ² = 17.91, df = 5, P = 0.003), indicating that the extent of staining or microbial recovery varied according to tissue position. In grafted plants injected in the scion, incidence did not differ significantly among treatments (Wald χ² = 1.39, df = 2, P = 0.500) or sampling positions (Wald χ² = 1.39, df = 1, P = 0.239), and pairwise comparisons confirmed the absence of significant differences among dye, M71, and SC1. For frequency, the treatment effect was marginal but not significant (Wald χ² = 5.36, df = 2, P = 0.069), whereas sampling position had a strong effect (Wald χ² = 32.04, df = 2, P < 0.001). SC1 tended to show higher frequency than the dye tracer, although this difference was not significant after FDR correction (P = 0.063). The position effect was mainly associated with higher recovery in the scion region.

### Spatial-temporal colonization of Pseudomonas chlororaphis M71 and Trichoderma atroviride SC1 in grapevine rooted cuttings

The persistence and spatial distribution of M71 and SC1 were assessed in rooted grapevine cuttings at 14, 28, and 56 dpi (Figure 3). The microorganisms were never recovered from SPSS-injected control plants.

**Figure 3.**
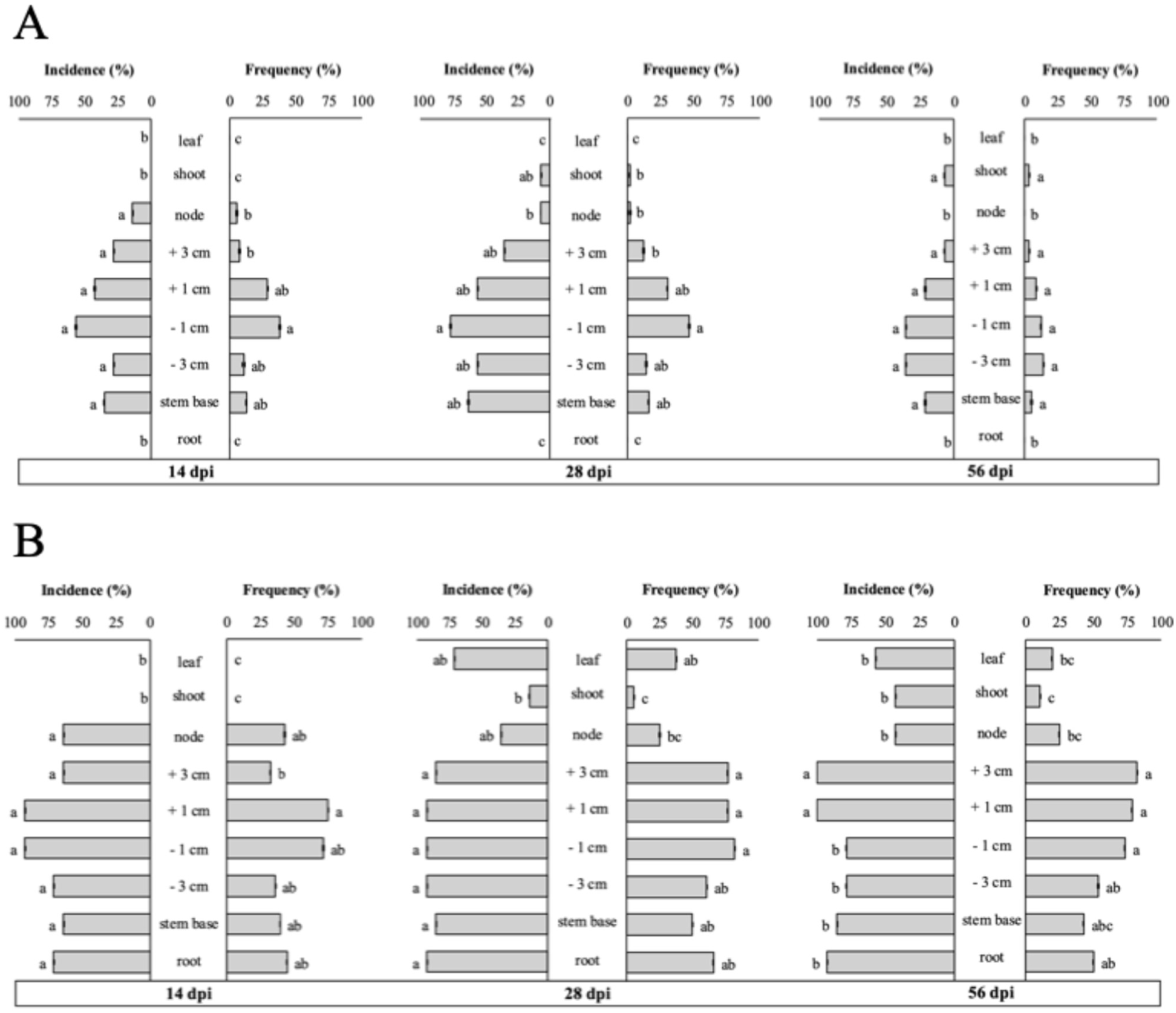
Incidence and frequency of Pseudomonas chlororaphis M71 (A) and Trichoderma atroviride SC1 (B), at 14, 28, and 56 days post-injection (dpi) in rooted grapevine cuttings. Colonization was assessed at various positions along the plant axis; + and - values indicate sampling positions above and below the injection site, respectively. At each time point, different letters indicate significant differences among positions according post-hoc test (P < 0.05). Error bar indicate standard error.

Following stem injection, M71 established in rooted cuttings and remained detectable over time, but its distribution was mainly restricted to stem tissues close to the injection site (Figure 3A). At 14 dpi, M71 was recovered from positions above and below the injection point, including +1, +3, -1, and -3 cm, as well as from the stem base and node, but not in leaves, shoots and roots. Incidence did not differ significantly among colonized positions at this time point (FDR-adjusted pairwise comparisons were non-significant). Frequency was highest at -1 cm, suggesting more intense local colonization close to the injection site. At 28 dpi, M71 was still recovered from several stem positions, confirming persistence for at least one month. Colonization was highest at -1 cm, with lower or intermediate values at +1, +3, -3 cm, the stem base, and the node. Pairwise comparisons for incidence showed a significant difference only between -1 cm and the node (P = 0.0189). Similar behavior is noted for frequency, with differences between -1 cm and shoot, node and + 3 cm. At 56 dpi, M71 remained detectable and its distribution remained localized and mainly associated with stem positions close to the injection site, whereas roots and leaves were never colonized. No significant differences among colonized positions were detected for incidence and frequency at this time point (all post-hoc comparisons were non-significant). Overall, M71 persisted after stem injection but showed limited systemic distribution, with little evidence of recovery from roots or distal aerial tissues.

Compared with M71, SC1 showed a broader and more persistent colonization pattern. At 14 dpi, SC1 was already detected in several stem positions and in roots, whereas distal aerial organs (shoot and leaf), were not colonized. Incidence was high in stem and root tissues, especially near the injection site, but no significant differences among positions were detected after post-hoc correction. Frequency, however, indicated greater colonization close to the injection site, particularly at +1 cm, with lower values in distal aerial tissues. At 28 dpi, SC1 showed its widest distribution. Incidence was high in most stem positions and in roots, whereas the shoot was the least colonized position. Post-hoc comparisons showed that shoot incidence was significantly lower than roots (P = 0.0207), stem base (P = 0.0255), +3 cm (P = 0.0207), -3 cm (P = 0.0207), -1 cm (P = 0.0207), and +1 cm (P = 0.0207). The node and leaf showed intermediate incidence and did not differ significantly from the other positions after correction. Frequency showed a similar spatial pattern, with higher values in roots, basal stem tissues, and positions close to the injection site, and lower values in distal aerial tissues. At 56 dpi, SC1 remained widely detectable. Incidence was relatively higher above the injection site and in the model, several position coefficients differed significantly from the + 1 and + 3 cm above the infection site. Frequency also showed a similar pattern, with relatively lower colonization only above the node. These data indicate that SC1 persisted and was recovered from broader range of positions in the plant than M71, although colonization intensity remained position-dependent.

Quantitative recovery from wood tissue at 28 dpi further highlighted differences between the two microorganisms. M71 reached 1.9 × 10⁵ CFU g⁻¹ at 1 cm above the injection point and 1.2 × 10⁷ CFU g⁻¹ at 1 cm below the injection point. SC1 was recovered at lower concentrations, reaching 1.3 × 10³ CFU g⁻¹ at 1 cm above and 1.2 × 10⁴ CFU g⁻¹ at 1 cm below the injection point.

Overall, both microorganisms persisted in rooted grapevine cuttings after stem injection, but their spatial-temporal patterns differed. M71 showed higher local recovery near the injection site but remained mainly confined to adjacent stem tissues, with limited or no movement to roots and distal aerial organs. SC1, although recovered at lower CFU levels from wood tissue, showed broader incidence and frequency across the plant, including roots and basal stem tissues. These results indicate that SC1 behaved as a more extensive endophytic colonizer than M71, whereas M71 showed a more localized colonization pattern under the conditions tested.

### Spatial-temporal colonization of Pseudomonas chlororaphis M71 and Trichoderma atroviride SC1 in grapevine grafted plants

The spatial-temporal colonization of M71 and SC1 was assessed in grafted grapevine plants after injection either in the rootstock or in the scion to understand the effect of the graft on colonization patterns. M71 and SC1 were never recovered from SPSS-injected control plants, confirming also in this case the absence of contamination.

When M71 was injected into the rootstock, recovery at 28 dpi was mainly concentrated around the injection area with limited detection toward the graft region, scion, or aerial organs (Figure 4A). Incidence was highest at +3 cm above the injection site (P = 0.0477) and other positions showed intermediate values. Frequency showed a similar localized pattern, with the highest recovery at -3 cm. By 56 dpi, M71 recovery after rootstock injection had markedly declined and was restricted to rootstock tissues close to the injection point. Incidence was significantly higher at -3 cm and +3 cm than at more distant positions and frequency confirmed this pattern. M71 was not recovered from leaves at either time point.

**Figure 4.**
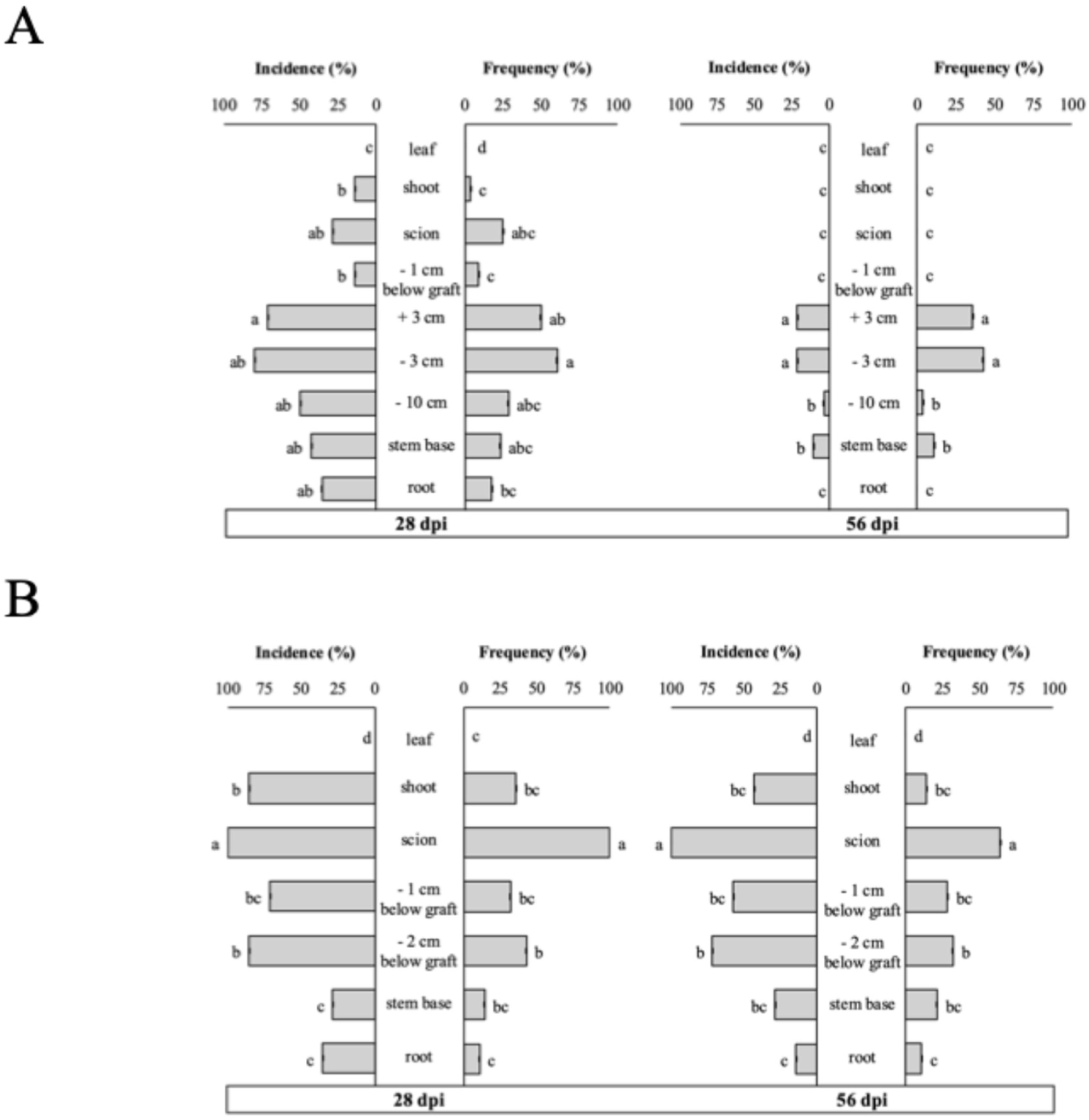
Incidence and frequency of Pseudomonas chlororaphis M71 at 28 and 56 days post-injection (dpi) in grafted grapevine plants injected in the rootstock (A) and in the scion (B). Colonization was assessed at different positions along the plant axis relative to the injection site; + and - values without specifications indicate sampling positions above and below the injection site, respectively. At each time point, different letters indicate significant differences among positions according post-hoc test (P < 0.05). Error bar indicate standard error.

When M71 was injected into the scion, colonization was more concentrated in the scion region and remained highly detectable over time (Figure 4B). At 28 dpi, incidence was highest in the scion, followed by the shoot and the region below the graft union. Frequency showed the clearest localization in the scion. The scion position had the highest recovery and differed significantly from roots (P < 0.0001), the stem base (P < 0.0001), 2 cm below the graft union (P = 0.0068), 1 cm below the graft union (P = 0.0004), and the shoot (P = 0.0011). At 56 dpi, M71 was still recovered mainly from the scion and nearby tissues, although overall recovery was generally lower than at 28 dpi. Incidence remained highest in the scion and frequency again confirmed the highest recovery in the scion. M71 did not migrate in the leaf at any time point.

Overall, on grafted plants the colonization pattern of M71 differed according to the injection site, but in both cases remained restricted to the tissues immediately surrounding the injection point and slightly declined between 28 and 56 dpi. Comparing the two injection sites, M71 displayed slightly different trends. In rootstock-injected plants, M71 showed a marked decline in both incidence and frequency between 28 and 56 dpi, a pattern that was less strong in scion-injected plants, where presence appeared more stable over the same period.

When SC1 was injected into the rootstock, colonization was already well established at 28 dpi and persisted until 56 dpi (Figure 5A). At 28 dpi, incidence was highest at +3 cm above the injection site, although most rootstock and basal positions were not clearly separated statistically. Incidence was significantly lower in the scion and leaf compared with the most colonized region (P = 0.0247), indicating limited upward movement into distal scion-derived tissues. Frequency had the highest recovery at -3 cm and high values also in the stem base and at +3 cm. By 56 dpi, SC1 was recovered from all sampling positions, with incidence highest at -3 cm. The shoot showed the lowest incidence. Frequency confirmed a similar pattern, with the highest recovery at +3 cm, -3 cm, and below the graft region, and significantly lower values in the shoot compared with -3 cm (P = 0.0049), +3 cm (P = 0.0008), and below the graft region (P = 0.0049). Thus, after rootstock injection, SC1 persisted mainly in rootstock and basal tissues, with partial movement toward the graft region but limited colonization of distal aerial tissues.

**Figure 5.**
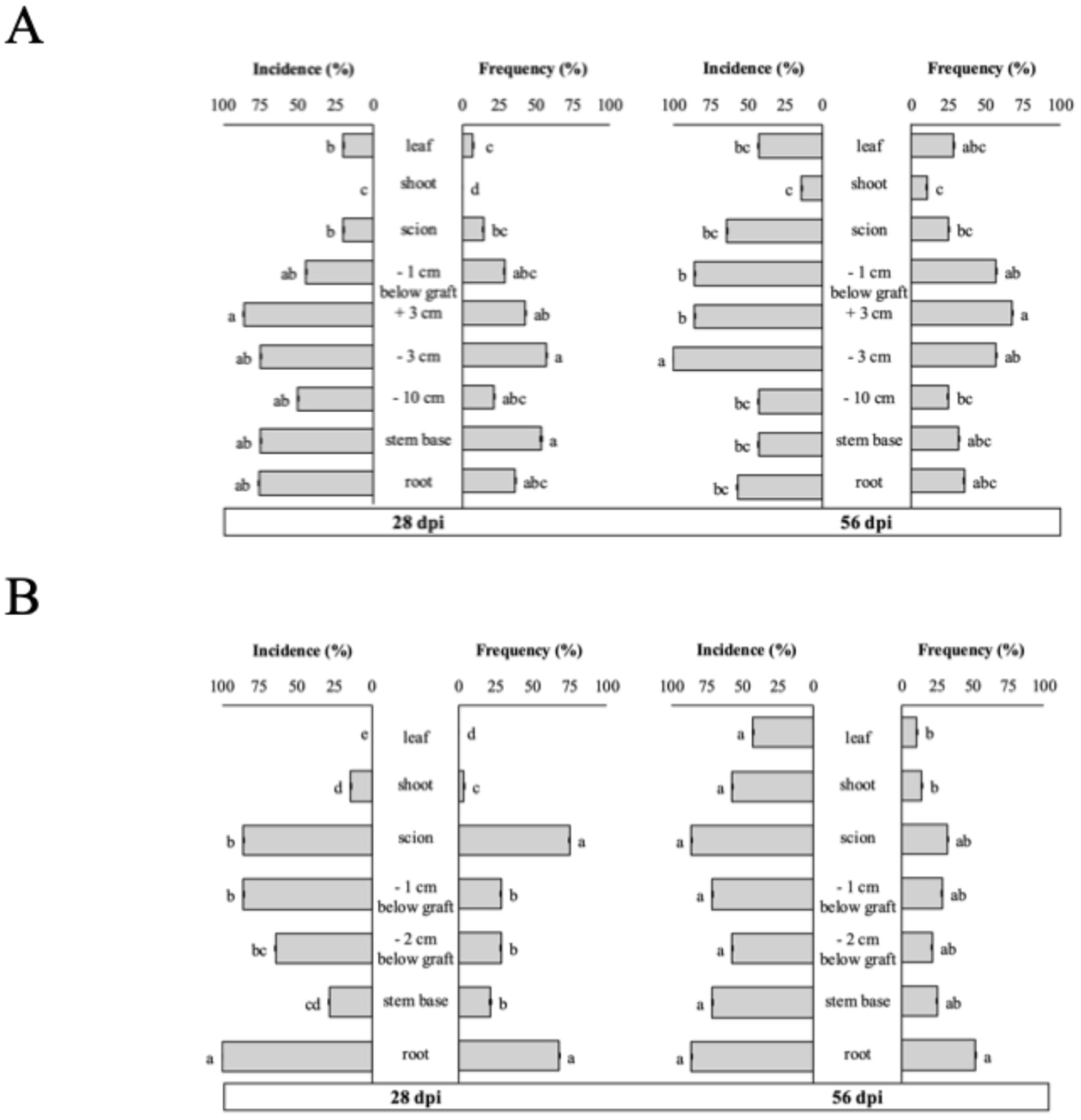
Incidence and frequency of Trichoderma atroviride SC1 at 28 and 56 days post-injection (dpi) in grafted grapevine plants injected in the rootstock (A) and in the scion (B). Colonization was assessed at different positions along the plant axis relative to the injection site; + and - values without specifications indicate sampling positions above and below the injection site, respectively. At each time point, different letters indicate significant differences among positions according post-hoc test (P < 0.05). Error bar indicate standard error.

When SC1 was injected into the scion, it showed a broader distribution across the grafted plant (Figure 5B). At 28 dpi, incidence was highest in the scion, roots, and at 1 cm below the graft union. Incidence in the scion and at 1 cm below the graft union was significantly higher than in the trunk base (P = 0.0147 for both) and shoot (P = 0.0092 for both), while the position 2 cm below the graft union showed higher incidence than the shoot (P = 0.0230). Frequency was highest in the roots and scion, which were statistically comparable. At 56 dpi, SC1 was recovered from all assessed regions, including the leaf. Incidence no longer differed significantly among positions after post-hoc correction, indicating widespread presence throughout the plant. Frequency, however, remained position-dependent, with the highest recovery in roots. Root frequency was significantly higher than in the shoot (P = 0.0215) and leaf (P = 0.0081), whereas the other woody tissues showed intermediate values. At 28 dpi SC1 was not found in the leaf, while at 56 dpi was present also in the leaf at comparable concentration to the other sampling sites

Overall, SC1 established and persisted in grafted plants after both rootstock and scion injection. After rootstock injection, colonization remained strongest in the rootstock and basal tissues, with partial movement across the graft region and reduced recovery in shoot and leaf tissues. After scion injection, SC1 showed broader recovery across the grafted plant, including recovery from roots and distal aerial tissues by 56 dpi. However, incidence and frequency did not always increase in parallel: SC1 could be detected across many positions, but local colonization intensity remained highest in woody and basal tissues and lower in green aerial organs.

### In planta visualization of Pseudomonas chlororaphis M71

To confirm the presence and tissue localization of M71 after stem injection, the strain was fluorescently labelled with a green fluorescent protein reporter through chromosomal insertion using a triparental mating approach. The transformed strain retained the main phenotypic and functional traits of the wild-type strain, including colony morphology, growth behavior, phenazine production, and antagonistic activity against *Phaeomoniella chlamydospora* and *Neofusicoccum parvum*. Antagonistic activity was assessed through dual confrontation assays on PDA at 26 °C, using agar-plug mycelium as the fungal inoculum and a 25 µL spot of bacterial suspension (1×10⁸ CFU mL⁻¹). Inhibition rates were comparable between the wild-type and the transformed strain: the wild-type inhibited *P. chlamydospora* by 59% and *N. parvum* by 48%, while the transformed strain achieved inhibition rates of 59% and 46%, respectively, with no statistically significant differences between the two strains.

At 28 days post-injection, GFP-labelled M71 was successfully detected in transverse wood sections of rooted grapevine cuttings (Figure 6). Fluorescent bacterial cells were observed inside the wood tissues both above and below the injection site. The signal was consistently associated with vascular tissues and was mainly localized within xylem vessels, indicating that M71 colonization occurred preferentially through the xylem. No clear GFP signal attributable to M71 was observed in phloem tissues, adjacent parenchyma, or pith in all observed samples.

**Figure 6.**
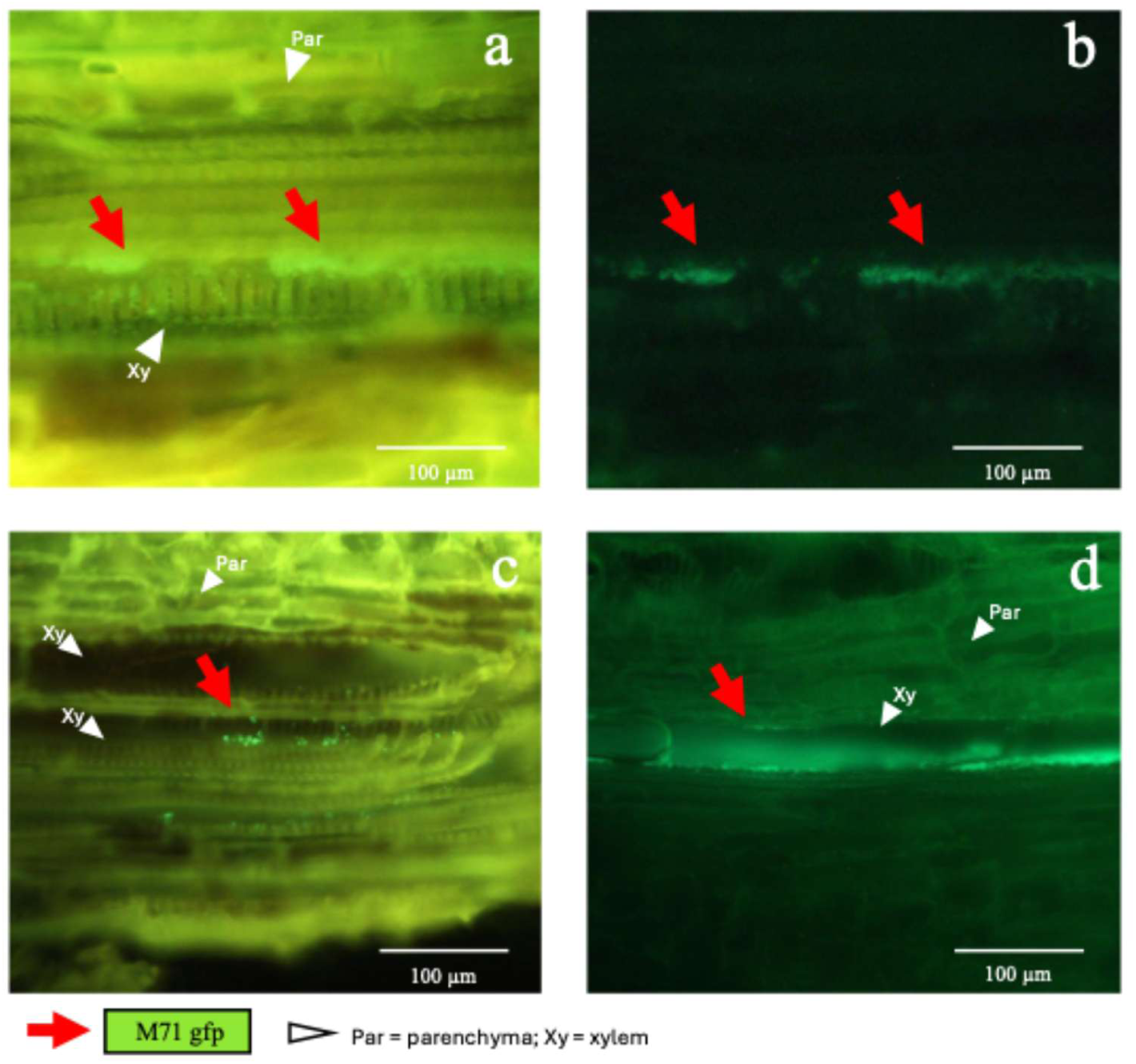
Fluorescence microscopy images of grapevine wood sections showing GFP-labelled Pseudomonas chlororaphis M71 inside xylem vessels. In the upper row, the same tissue section was observed using different fluorescence DAPI filter sets: (A) excitation 450-490 nm, B-2A filter and (B) excitation 420-490 nm, B-3A filter. In the lower row, different images show M71 cells with B-2A filter (C), as well as with B-3A filter (D).

The presence of viable GFP-labelled M71 was further confirmed by plating corresponding wood sections on rifampicin-amended medium. Fluorescent colonies were recovered from M71-injected cuttings and showed the same fluorescence pattern under microscopy, confirming that the signal detected *in planta* corresponded to viable labelled bacteria. No alteration of grapevine internal tissues was observed in M71 injected samples. No fluorescent colonies were obtained from all samples of SPSS-injected control cuttings, confirming the absence of contamination and background recovery. The DAPI filter allowed to distinguish plant tissue autofluorescence from bacterial GFP fluorescence. Wood sections were examined under three excitation filters: B-3A (420–490 nm), which revealed only GFP-labelled bacterial cells (Figure 6); B-2A (450–490 nm), which showed both bacterial and wood tissue signals (Figure 6); and a third filter (340–380 nm), which detected only plant tissue autofluorescence with no bacterial signal (Image 1, supplemental material). This confirms that the GFP signal observed under the B-3A filter was exclusively related to M71 cells and not to background autofluorescence of wood tissues.

All these observations indicate that GFP-labelled M71 survived and colonized grapevine wood tissues after stem injection. Its localization within vascular tissues, particularly xylem vessels, supports the colonization assay results and suggests that M71 dispersal and persistence in planta are mainly associated with the xylem system.

### In planta visualization of Trichoderma atroviride SC1

To confirm the presence and tissue localization of SC1 after stem injection, rooted grapevine cuttings were examined by scanning electron microscopy at 28 days post-injection. Hyphae were detected in all wood samples from SC1-injected plants, particularly within vascular tissues (Figure 7). These fungal structures were observed mainly inside xylem vessels, indicating that SC1 colonized internal grapevine tissues rather than remaining confined to the stem surface. Viable SC1 was consistently recovered from the corresponding wood sections plated on semi-selective medium, confirming that the fungal structures observed by microscopy were associated with living SC1 colonization. No alteration of grapevine internal tissues was observed in SC1 injected samples. In contrast, neither hyphae nor SC1 colonies were detected in SPSS-injected control plants, confirming the absence of contamination and the specificity of the observations. Therefore, SEM analysis supports the hypothesis that SC1 survived and established within grapevine wood tissues after stem injection and indicates that SC1 can behave as an internal colonizer of rooted grapevine cuttings.

**Figure 7.**
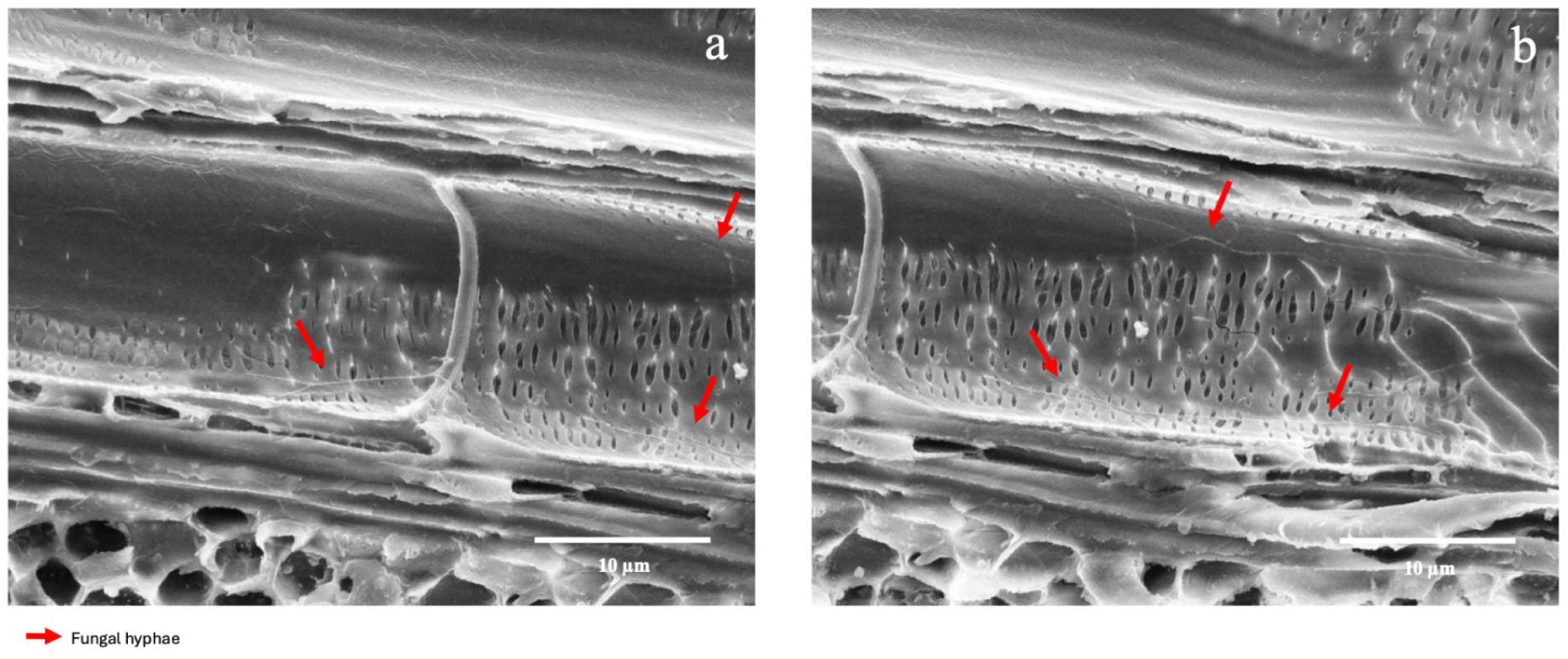
Scanning electron microscopy image of hyphae ascribed to Trichoderma atroviride SC1 inside a xylem vessel (red arrows) of rooted cutting 28 days post injection with SC1 conidial suspension (1×10^6^ conidia mL⁻¹). Hyphae were absent from all samples injected with SPSS.

## Discussion

This study assessed whether two beneficial microorganisms, *Pseudomonas chlororaphis* M71 and *Trichoderma atroviride* SC1, can be introduced into grapevine tissues by stem injection, persist internally, and be recovered from rooted and grafted plants without causing visible phytotoxic effects.

The stem injection method was effective and reproducible in all experimental systems. The dye tracer confirmed that the injected suspension entered the vascular system and moved mainly through the xylem, without inducing visible phytotoxicity.

At 48 h post-injection, dye movement was generally broader along the plant axis than microbial recovery, indicating that passive fluid transport occurred more rapidly than microbial colonization. However, dye staining was laterally restricted within each section, being mostly confined to xylem vessels adjacent to the injection pathway. By contrast, when M71 or SC1 were recovered from the same sampled position, they were often detected in a wider proportion of tissue fragments. This suggests that the dye mainly reflected immediate passive transport within xylem, whereas microbial recovery likely reflected both passive transport and subsequent survival or growth within colonized tissues.

The early distribution of both microorganisms was associated with vascular transport, but their subsequent behavior differed. M71 showed a distribution pattern more closely linked to the injection pathway and remained largely restricted to tissues close to the point of inoculation. This is consistent with a bacterium mainly relying on passive transport through sap flow and with limited ability to actively explore surrounding tissues. In contrast, SC1 showed broader colonization over time and was frequently recovered from positions that were not strongly marked by the dye. This suggests that SC1 was not limited to passive displacement through the xylem but was also able to establish and extend within internal tissues, likely through hyphal growth. These observations are consistent with the known ability of endophytes to colonize internal tissues without causing visible damage (13), and with the capacity of microorganisms to use wounds or natural openings as entry points (2, 14, 15).

In rooted cuttings, both microorganisms persisted after injection, but their spatial-temporal patterns were clearly different. M71 remained mainly localized around the injection site and was only weakly or never recovered from distal aerial tissues or roots. Although M71 reached high local populations in wood tissue at 28 dpi, its distribution remained spatially restricted and declined over time. This behavior may reflect dependence on xylem-mediated movement and limited adaptation to grapevine woody tissues. Similar constraints have been reported for introduced biocontrol agents, whose populations often decline after inoculation because of host genotype, inoculation method, microbial competition, or limited adaptation to internal plant niches (20, 28). The restricted persistence of M71 may also reflect strain-host specificity, as other *Pseudomonas* strains have shown different persistence capacities in grapevine or woody plant systems (36, 47). Alternatively, as a non-pathogenic bacterium, M71 may lack traits required to overcome host barriers or maintain long-term proliferation within woody tissues, in line with observations of James et al. (2002) (48).

In contrast, SC1 showed a broader and more persistent colonization pattern than M71. In rooted cuttings, SC1 was recovered from stem tissues and roots from the early sampling points and remained detectable up to 56 dpi. This agrees with the known ecological behavior of *Trichoderma* spp., which are well adapted to the rhizosphere and can colonize plant tissues as endophytes (49). The recovery of SC1 from roots, despite stem injection, is particularly relevant because dye movement did not indicate strong passive transport toward the root system. This suggests that SC1 movement was not solely driven by xylem flow but may have involved active growth toward favorable niches or host-derived signals, consistent with the role of microbial chemical signaling (50). The persistence of SC1 without visible tissue alteration also supports a compatible interaction with grapevine tissues, potentially involving modulation of host responses, as reported for *Trichoderma*-plant interactions (51, 52).

The quantitative isolation data further highlighted the different behavior of the two microorganisms. M71 reached higher CFU values than SC1 at 28 dpi, especially below the injection point, but this did not translate into broader systemic colonization. SC1, although recovered at lower CFU levels, showed greater spatial distribution and persistence. This distinction most probably reflects the cell load at the inoculation but also suggests that high local population density does not necessarily correspond to systemic spread. M71 behaved as a strong local colonizer of specific vascular compartments, whereas SC1 showed lower local density but greater ability to occupy multiple plant regions over time.

The grafted plant experiments showed that the graft union influenced microbial distribution, but did not completely prevent translocation. Both microorganisms were recovered across grafted tissues, indicating that vascular continuity in young grafted plants was sufficient to allow some microbial movement despite recent callus formation, new vascular differentiation, and partial vessel obstruction typical of grafted nursery material (53, 54). However, the two microorganisms responded differently to the grafted structure. M71 remained mostly restricted to the region surrounding the injection point, regardless of whether it was injected into the rootstock or scion. After rootstock injection, its recovery declined strongly between 28 and 56 dpi, whereas after scion injection it persisted more consistently, although mainly in the scion and adjacent tissues. This may reflect differences in tissue physiology between rootstock and scion, including local vascular activity and proximity to actively growing aerial organs. SC1 displayed greater capacity to colonize grafted plants rather than rooted cuttings. After rootstock injection, it persisted mainly in rootstock and basal tissues, with partial movement toward the graft union and limited colonization of shoot and leaf tissues. After scion injection, SC1 was recovered from both aerial tissues and basal/root tissues, and by 56 dpi it was detected across all assessed regions, including leaves. These results indicate that the graft union did not prevent SC1 movement, although colonization intensity remained higher in woody and basal tissues than in distal green organs. The delayed and position-dependent recovery suggests that SC1 may require an establishment phase before extending through structurally distinct tissues across the graft union. This behavior is consistent with the adaptive plasticity of *Trichoderma* morphology and their ability to colonize different plant compartments (55, 56).

Direct microscopy supported the isolation data and confirmed the internal localization of both microorganisms. GFP-labelled M71 was observed inside grapevine wood tissues, mainly within xylem vessels, and viable fluorescent colonies were recovered from corresponding wood sections. Similarly, SEM observations showed SC1 hyphae inside xylem vessels, and viable SC1 was recovered from the same tissues. No corresponding structures or colonies were observed in SPSS-treated controls. These observations demonstrate that both microorganisms colonized internal plant tissues rather than remaining on the external stem surface. The preferential localization of M71 in xylem vessels is consistent with a strategy based on passive vascular transport. Xylem vessels can facilitate longitudinal movement without requiring active exploration of parenchyma or pith tissues (15, 57). For SC1, the presence of hyphae in xylem vessels suggests that these conduits may function as dispersal pathways, as proposed for endophytic fungi by Boddy (1994) (58), while hyphal growth may allow subsequent colonization of adjacent tissues. Importantly, no tissue disruption or vessel occlusion was observed, supporting an endophytic rather than pathogenic interaction (59).

The contrasting behavior of M71 and SC1 reflects their different biological traits. M71, as a bacterium, appeared to rely mainly on xylem flow and showed efficient but localized colonization. Its persistence may be useful where transient or spatially targeted activity is desired, although disease protection after injection remains to be tested. SC1, as a filamentous fungus, combined initial passive movement with active internal growth, resulting in broader and more persistent colonization. This pattern supports the potential of SC1 as a systemic or semi-systemic endophytic biocontrol agent in grapevine woody tissues. Field studies have shown that *Trichoderma* spp. applications to pruning wounds can reduce Grapevine Leaf Stripe Disease (GLSD, within Esca complex disease)-related symptoms and that SC1 can be recovered several months after application from grapevine wood (60), supporting the relevance of the colonization behavior observed here under controlled conditions.

These findings may have practical implications for grapevine biocontrol. Introducing beneficial microorganisms into internal tissues during nursery production or early plant development could help establish protective endophytic populations before pathogen infection. This is particularly relevant for grafting, which represents a critical stage for pathogen entry and for the shaping of the grapevine endophytic community (61). Previous work has shown that endophytic bacteria can reduce infections in nursery plants, as demonstrated by Álvarez-Pérez et al. (2017) (62), and the use of SC1 to protect rootstocks against Esca Complex-associated fungi during grafting has already been demonstrated by Pertot et al. (2016) (63). However, persistence under field conditions and the contribution of these microorganisms to long-term plant protection remain to be clarified (64). Moreover, although endophytes can improve plant health, immune responses triggered by endophytes or their metabolites may occasionally affect tissue integrity or plant fitness, especially in young plants (65, 66).

The ability of M71 and SC1 to occupy different internal niches suggests that their use could be complementary. M71 may provide localized and rapid activity within vascular tissues, whereas SC1 may offer longer-term establishment and broader tissue colonization. Combined or sequential application strategies could therefore be explored, particularly during propagation or early field establishment. This may be relevant for vascular and woody diseases in which the structure of the endophytic microbiome influences disease progression, including grapevine trunk diseases, Flavescence dorée, Olive Quick Decline Syndrome, and Pierce’s disease (67, 68, 69, 70). Pathogens such as *Xylella fastidiosa* and phytoplasmas can strongly alter endophytic communities (71, 72), and grapevines affected by trunk diseases often show reduced populations of potentially beneficial endophytes such as *Pseudomonas* species. in diseased tissues (37). Moreover, initial evidence of a potential endotherapeutic effect of SC1 against *Gnomoniopsis castanea* in chestnut has been observed (73). These observations support the hypothesis that targeted manipulation of the endophytic microbiome could contribute to disease suppression or plant recovery, although this hypothesis requires direct testing in pathogen-challenge and field experiments.

Although strain recovery was supported by rifampicin selection for M71, the isolation and identification protocol described by Savazzini et al. (2008) for SC1 (45), and microscopy observations for both M71 and SC1 and therefore identification based on these methods was considered sufficiently robust to confirm the identity of the two microorganisms, a limitation of the study is the absence of molecular confirmation for all recovered isolates.

In conclusion, this study demonstrates that stem injection can be used to introduce beneficial microorganisms into grapevine tissues and to study their internal movement, persistence, and tissue localization. M71 and SC1 both survived after injection, but they followed different colonization patterns. M71 remained primarily xylem-associated and localized to the site of injection, whereas SC1 had broader and more persistent internal colonization. The graft union did not prevent microbial movement, although it partially influenced the extent and direction of colonization. Future work should include transcriptomic analyses to clarify host-microbe interactions and field trials to determine whether the persistence and distribution observed in potted plants translate into stable colonization and disease protection under vineyard conditions.

## Acknowledgments

The authors thank David Baldo for technical assistance with microscopy, Oscar Giovannini for support with greenhouse plant maintenance, and Carmela Sicher, Maricarmen Costigliola, and Laila Cattane for assistance with laboratory activities. Special thanks to Stefano Di Marco for his encouragement and support at the beginning of this study. The authors used ChatGPT by OpenAI (GPT-5.5) only for English-language editing and improvement of clarity and readability. All scientific content, data analysis, interpretation, and conclusions were produced and verified by the authors.

## Author Contributions

Greta Brussi and Ilaria Pertot conceived the study and designed the experiments. Greta Brussi performed the stem-injection experiments, microbial isolations and colonization assays. Greta Brussi and Alessio Martini generated and characterized the GFP-labeled M71 strain. Greta Brussi performed fluorescence microscopy. Greta Brussi and Claudio Ratti performed scanning electron microscopy. Greta Brussi analyzed the data, prepared the Figures and wrote the original draft. Ilaria Pertot and Gerardo Puopolo supervised the work. Ilaria Pertot and Laura Mugnai reviewed and edited the manuscript.

